# Rapid and widespread transcriptional plasticity underlies host shift in a specialist herbivore

**DOI:** 10.1101/2024.10.15.618594

**Authors:** W. Nate Collison, Syed Adeel Zafar, Simon C. Groen, Paul D. Nabity

## Abstract

Insect herbivores employ many adaptive mechanisms to successfully colonize host plants. Transcriptional plasticity is one mechanism that enables broad host use in polyphagous insect herbivores; however, whether specialist herbivores use transcriptional plasticity or undergo epigenome modifications to cope with host shifts remains largely unexplored. We measured the transcriptome and DNA methylome of the specialist aphid herbivore, *Daktulosphaira vitifoliae*, after transfer to new hosts using RNA-seq and whole-genome bisulfite sequencing. We found the transcriptome of *D. vitifoliae* is highly plastic and responsive to host genotype – 3,706 genes (28% of expressed genes) are significantly differentially expressed in at least one contrast comparing host genotypes. Most host-responsive genes (2,667 genes) were upregulated when feeding on the non-native host *Vitis vinifera* and enriched for genes encoding predicted secretory effectors, detoxification enzymes, and cuticle- and chitin-associated proteins. We also found gene co-expression modules that correlated with variation in fitness suggesting plasticity in transcriptional networks contributes to *D. vitifoliae* adaptation to host plants. We also identified a previously undescribed family of proline-rich secretory proteins conserved across the Aphidomorpha infraorder of insects. Our analysis of DNA methylation revealed a negative correlation between gene-level CpG methylation and host-induced transcriptional plasticity, suggesting a lack of gene body methylation may enable transcriptional plasticity when insects move to new host environments. Together, these results provide new insight into the molecular mechanisms of host use in insect herbivores.

## Introduction

Phenotypic plasticity, or the expression of different phenotypes by the same genotype, can allow organisms to flexibly respond to environmental changes, and has thus been widely proposed to contribute to the evolution of species (Ancel, 2000; Pigliucci & Murren, 2003; Ghalambor et al., 2007; Murren et al., 2015). However, the extent to which plasticity alters adaptive evolution remains an open question (Vinton et al., 2022). Plant-herbivore interactions are particularly useful systems for understanding the mechanisms of plasticity and adaptation, given their ecological diversity (Futuyma & Agrawal, 2009; Forister et al., 2015), ability to rapidly adapt to new environments (Garnas, 2018), and dramatic examples of environment-induced phenotypic variation (Whitman & Agrawal, 2009).

Insect herbivores are among the most species-rich lineages of metazoans, and this diversity is largely driven by adaptations to specific host plants (Simon et al., 2015). These adaptations function as a mode of ecological specialization that induces trade-offs in fitness as herbivorous insects transition from ancestral to alternative hosts (García-Robledo & Horvitz, 2012). Support for this hypothesis comes from the fact that most insect herbivores are monophagous or oligophagous (i.e. specialists), feeding on one or very few species of closely related hosts (Bernays & Graham, 1988; Futuyma & Agrawal, 2009; Forister et al., 2015). For polyphagous, generalist herbivores, compatibility across a breadth of divergent hosts is hypothesized to be maintained by a high degree of genomic flexibility (Sexton et al., 2017; Birnbaum & Abbot, 2020). Indeed, the genomes of generalist insect herbivores contain extraordinary expansions of chemoreception and detoxification genes (W. Xu et al., 2016; Grbić et al., 2011; Pearce et al., 2017), which enable host finding and feeding on plants. In contrast, specialist herbivores appear to lack such expansions of chemoreception and detoxification genes (Peláez et al., 2023). Also, there is increasing evidence that insect herbivores, both generalists and specialists, rely on expanded families of secretory effector proteins to directly manipulate host plant biology during feeding (Zhao et al., 2015; Mathers et al., 2017; Rispe et al., 2020; Korgaonkar et al., 2021).

Transcriptional plasticity of expanded gene families has been linked to fitness variation in generalist insect herbivores during interactions with divergent host plants (Mathers et al., 2017; Snoeck et al., 2018), suggesting transcriptional plasticity plays an adaptive role in these insects. Most studies that investigated transcriptional responses of insects to different host plants used generalists, as these insects typically represent major agricultural pests (Grbić et al., 2011; Mathers et al., 2017; Pearce et al., 2017). Transcriptional plasticity is hypothesized to be reduced in specialist herbivores relative to generalists (Birnbaum & Abbot, 2020), given host-specific evolution, i.e. reduced diet breadth. Indeed, the few studies that measured transcriptomic responses to host transfer in both specialist and generalist insects found overall reduced transcriptional plasticity in specialists (Govind et al., 2010; Roy et al., 2016; Schweizer et al., 2017). From an evolutionary perspective, specialization and the reduced capacity for plasticity are expected to come at a cost, limiting adaptability of species amidst rapid environmental change (Futuyma & Moreno, 1988; Price et al., 2003; Forister et al., 2012; Vamosi et al., 2014; Murren et al., 2015; Day et al., 2016).

Dynamic regulation of the epigenome is thought to be a central mechanism underlying rapid, environment-induced transcriptional plasticity (Duncan et al., 2014; Kilvitis et al., 2017; Duncan et al., 2022). DNA methylation of cytosines, the covalent addition of a methyl group to these residues, is a well-known epigenetic mark that maintains functional gene regulation in many eukaryotic organisms (Zemach et al., 2010; Schmitz et al., 2019). Cytosine methylation, hereafter DNA methylation, has been well-studied in some insect lineages, primarily social Hymenoptera for its role in mediating behavioral and phenotypic plasticity (Yan et al., 2014, 2015). Unlike in plants and mammals, DNA methylation in insects occurs at relatively low levels and is found primarily in gene body regions of actively expressed genes. Interestingly, some insects lack discernable levels of DNA methylation in their genomes, suggesting its role in gene regulation is dispensable (Bewick et al., 2016; Provataris et al., 2018). In insects with abundant DNA methylation the observed variation has been linked to developmental and phenotypic plasticity (Glastad et al., 2014, 2019). Early studies in honey bees (*Apis mellifera*) found differential DNA methylation coincides with differential gene expression (Kucharski et al., 2008; Li-Byarlay et al., 2013) whereas more recent studies did not (Harris et al., 2019; Marshall et al., 2019; Cardoso-Júnior et al., 2021). However, in other insects such as the green peach aphid (*Myzus persicae*), differential DNA methylation is strongly correlated with differential expression of sex-linked genes (Mathers et al., 2019), suggesting DNA methylation regulates transcriptional plasticity and the emergence of different phenotypes in some contexts, i.e. insect lineage or trait in question.

The degree to which phenotypic plasticity, and the underlying epigenetic and transcriptional plasticity, contribute to adaptation and evolution remains contentious (Pigliucci & Murren, 2003; Price et al., 2003; Ghalambor et al., 2007; Vinton et al., 2022). Plasticity may slow the rate of genetic evolution by shielding genotypes from the effects of natural selection, because one genotype expresses multiple phenotypes across environments (Ancel, 2000; Price et al., 2003; Paenke et al., 2007). Alternatively, plasticity that produces phenotypes which are favored by natural selection may later become genetically assimilated, a case in which plasticity results in adaptive evolution (Price et al., 2003; Paenke et al., 2007; Lande, 2009). Whether plasticity is adaptive depends on the strength and direction of selection. Plastic responses may move individuals towards a fitness optimum (adaptive), away from a fitness optimum (maladaptive), or have a neutral effect (Pigliucci & Murren, 2003; Ghalambor et al., 2007). A separate but related question is *to what extent does environment-induced plasticity lead to phenotypic adaptation via transgenerational inheritance of epigenetic marks*? The answer to this question similarly remains contentious, and likely dependent on the lineage or trait in question. The transfer of DNA methylation marks across generations in mammals seems to be rare, because almost all DNA methylation is erased then reestablished during embryogenesis (Morgan et al., 2005; H. J. Lee et al., 2014; S.-M. Lee & Surani, 2024). In insects, conversely, DNA methylation erasure during embryogenesis does not occur (Harris et al., 2019; X. Xu et al., 2019), and DNA methylation can be inherited across generations in *Nasonia* wasps (Wang et al., 2016) and honey bees (Yagound et al., 2020). Thus, whether inheritance of epigenetic marks can contribute to phenotypic adaptation in insects remains an open question.

In our study, we examined host-induced variation in fitness, gene expression, and DNA methylation in the specialist herbivore *Daktulosphaira vitifoliae* (Fitch). This gall-forming aphid-like insect, commonly known as grape phylloxera, is in the Phylloxeridae family (Aphidomorpha infraorder) of Hemipteran insects and feeds strictly on grapevines (species within the *Vitis* genus of flowering plants) (Granett et al., 2001; Metz et al., 2017). Although native to North America, *D. vitifoliae* has invaded grape growing regions across the globe, becoming one of the most infamous pests of grapevines (Rispe et al., 2020). The accidental introduction into France in the late 1800’s caused the ‘phylloxeric plague’, wiping out thousands of vineyards and threatening global grape production (Rispe et al., 2020). While agriculturally relevant, *D. vitifoliae* also provides a model study system for ecological genomics and epigenetics. Short generation times of asexually reproducing individuals allow for large clonal populations to be reared and removes any confounding effects of genetic background on transcriptional or epigenetic plasticity, which is a notable advantage in our study. The obligate nature of *D. vitifoliae* should also strengthen the ability to detect how host switching shapes trait plasticity, given the outcome is adapt or die. To explore these traits, we investigate the transcriptional and epigenetic basis of host use in *D*. *vitifoliae* as it feeds on different host genotypes of native and novel origin for a single generation. We test the hypothesis that plastic responses of the transcriptome mediate variation in *D. vitifoliae* fitness on different hosts, such that the direction of plasticity in transcriptional networks is positively correlated (adaptive) or negatively correlated (maladaptive) with fitness variation. We also analyzed DNA methylation in *D. vitifoliae* to characterize the DNA methylome and to assess the correlation between host-induced transcriptional plasticity and DNA methylation. Additionally, we analyzed DNA methylation in the progeny (eggs) of adults that fed on different host plants for one generation to assess whether feedings on different host plants led to inter-generational changes in DNA methylation. While we do not directly test for heritability of epigenetic marks, we test the hypothesis that environment-induced transcriptional plasticity leads to predictable changes in the epigenome of *D. vitifoliae* progeny after one generation in different host environments, providing new insight to the role of DNA methylation in phenotypic adaptation in insects.

## Materials and Methods

### Plant and insect rearing

A clonal line of *D. vitifoliae* was established from a single female leaf gall collected on *V. rupestris* (cv St. George) near Davis, California U.S.A., in 2021. The clonal population was maintained for successive generations on non-native *V. vinifera* cv Cabernet Sauvignon 33 (Cab33) that were grafted (*Vitis* genotype 1616 and 3309 rootstocks) and self-rooted in UC Riverside greenhouses in Riverside, CA (27 °C +/-3 °C) until experimentation during 2022-2023.

We used eight different grapevine genotypes as hosts for *D. vitifoliae* in our experiments, all of which were clonally propagated from cuttings purchased as rooted, overwintered 1-year-old plants from Inland Desert Nursery (Benton City, WA). Genotypes included self-rooted rootstock hybrids of *V. berlandieri* x *V. rupestris* (99R, 1103P), *V. berlandieri* x *V. riparia* (SO4, 5BB), and *V. riparia* x *V. rupestris* (101-14, SW, 3309). All plants were potted in 11.3 liter round pots with ‘UC Mix 3’ soil (https://agops.ucr.edu/soil-mixing#uc-soil-mix-iii) and grown in a greenhouse set at 24:27 °C ± 3 °C (night:day) with a 14h photoperiod. After leaf bud swelling (8 days after potting), half of the plants for each genotype were infested. The remaining plants were potted and infested similarly 6 days later. Plantings were replicated through time to optimize sample collection effort given the time required to sort and count insects.

### Host transfer and insect performance assay

For experimental infestation of new host plants (host transfer), galled leaves from the colony on Cab33 (hereafter ‘G0’, for generation 0) were excised and placed on the branch nodes of new host plants. Infestation was monitored every 12 hours to note hatch and dispersal of 1^st^ nymphal instars (hereafter generation 1, ‘G1’) onto budding leaves. Galled leaves used for infestation were removed from branch nodes after two days to prevent further infestation.

To assess insect performance, ‘G1’ galls were collected 15 days post infestation (DPI). Galls were collected using a 4 mm diameter hole punch and carefully opened under a dissecting microscope to count eggs per gall. Each plant was sampled five different times, at 15, 17, 19, 21, and 23 DPI. Each time, the five largest galls from each plant were sampled, selecting galls from leaves that had not been previously sampled unless the largest remaining galls were on a previously sampled leaf. To determine if *D. vitifoliae* fitness varied significantly on different hosts, pairwise Wilcoxon rank-sum tests were performed (R v. 4.3.1).

### RNA isolation and sequencing

To quantify *D. vitifoliae* transcriptome plasticity during host transfer events, twenty G1 adults were collected from galls on unique plants and pooled for each sample. Samples were collected 15-23 DPI and placed in tubes on ice during collection, then immediately stored at -80 °C until RNA isolation. RNA was isolated from whole bodies using the RNeasy Mini Kit (Qiagen) following manufacturer’s instructions. RNA intactness and purity were verified with gel electrophoresis and spectrophotometry (260:280 > 2; 260:230 >1.7) prior to library preparation using NEBNext^®^ Ultra^™^ II Directional RNA Library Prep Kit for Illumina^®^ (New England Biolabs). RNA libraries were sequenced using an Illumina NovaSeq X system at the DNA Technologies and Expression Analysis Core of the UC Davis Genome Center, Davis, CA, with an average yield of 28.9 (± 3.29 s.d.) million paired-end reads per sample. Sequencing data were uploaded to the NCBI Sequence Read Archive (SRA) repository under Bioproject accession PRJNA1211173.

### RNA-seq data processing

Fastqc v. 0.11.9 (https://github.com/s-andrews/FastQC) was used to generate read and base quality statistics for raw and processed reads. Raw RNA-seq reads were quality trimmed and filtered using BBTools bbduk (v.38.95; https://jgi.doe.gov/data-and-tools/software-tools/bbtools/) with default settings plus options for kmer length (k=23) and hamming distance (hist=1). Trimmed and filtered reads were aligned to the reference genome (Rispe et al., 2020) using STAR v. 2.7.10b (Dobin et al., 2013) with default settings plus options for maximum intron length (--alignIntronMax 15000) and removing alignments that contained non-canonical junctions (--outFilterIntronMotifs RemoveNoncanonical). On average, 85.8% (± 1.49% s.d.) of reads uniquely mapped to the reference genome, yielding 6.80 Gigabases (± 0.74 s.d.) of read coverage (Table S1). Only uniquely mapped reads were used for downstream analysis.

### Differential gene expression analysis

Gene count matrices were generated using the --quantMode GeneCounts option in STAR and analyzed using edgeR (v.4.0.16) (Robinson et al., 2010). Low count genes were removed from downstream analysis using filterByExpr with default parameters. Linear model fitting of count data was performed using limma’s voom and lmFit functions (Law et al., 2014). Genes were considered differentially expressed based on the empirical Bayes method implemented using limma’s ‘treat’ function with a 1.5-fold-change filter and BH-adjusted p-value < 0.05. Genes that were differentially expressed between ‘G1’ Cab33 and ‘G0’ (initial *D. vitifoliae* colony on Cab33) were removed from downstream analysis of host-responsive genes, since these were differentially expressed across generations on the same host genotype. Fisher’s exact testing for gene functional group enrichment was performed with the R stats package (v.4.3.1). Functional groups were considered significantly enriched based on a Benjamini Hochberg (BH)-adjusted p-value cutoff of 0.05 and an odds ratio ≥2. Functional groups were defined based on published genome annotations (Rispe et al., 2020).

### Gene co-expression network analysis

Genes with 0.5 counts per million (CPM; about 10 raw gene counts) in at least four samples were included for network construction using WGCNA (Langfelder & Horvath, 2008). Gene count matrices were normalized using DESeq2’s vst (variance stabilizing transformation) function (v. 1.42.1). A signed gene co-expression network was constructed based on hierarchical clustering of dissimilarity in gene connectivity (1-TOM) with the soft power parameter (β) set to 16 to ensure the network had approximate scale-free topology. Genes were assigned to modules using the cutreeDynamic function with minimum module size = 20 and module detection sensitivity (deepSplit) = 2. Heat maps of module eigengenes and normalized gene expression values (z-scores of vst-transformed count data) were made using pheatmap v. 1.0.12 (https://github.com/raivokolde/pheatmap; rows were clustered based on Euclidean distance. Correlations between *D. vitifoliae* fitness and gene co-expression modules were tested by linear regression of average fitness on each host genotype and average module eigengene expression using the lm function in the R stats package v. 4.3.1 where mean fitness = β0 + β1 × mean eigengene + ɛ. Scatter plots of eigengene-fitness correlations were printed with results from the linear regression test using ggplot2 v. 3.5.0 (https://ggplot2.tidyverse.org/). Gene co-expression modules were tested for overrepresentation/enrichment of gene functional groups based on Fisher’s exact tests as described above.

### KEGG term enrichment analysis

eggNOG-mapper v. 2 (Cantalapiedra et al., 2021) was used to assign KEGG terms to the predicted proteome of *D. vitifoliae* based on precomputed orthology assignments. eggNOG-mapper assigned KEGG terms to 7,532 proteins out of 15,076 queried. To test for KEGG term enrichment for subsets of genes (e.g. differentially expressed genes vs all KEGG-annotated genes) we performed hypergeometric over-representation analysis using the ‘enrich’ function of the R package clusterProfiler v. 4.10.1 (G. Yu et al., 2012). KEGG terms were considered significantly enriched based on a BH-adjusted p-value cutoff of 0.05 and a gene count ≥4.

### Multiple Sequence Alignment and phylogenetic analysis

To determine sequence similarity and phylogenetic relationships of *D. vitifoliae* proteins and putative homologs in other species, we performed Multiple Sequence Alignment (MSA) using the Clustal Omega web application (https://www.ebi.ac.uk/jdispatcher/msa/clustalo; Sievers & Higgins, 2018). Alignment results were visualized in a sequence logo plot using the WebLogo web application (https://weblogo.threeplusone.com; Crooks et al., 2004). Phylogenetic trees were constructed from the Clustal Omega MSA output using the Interactive Tree of Life (iTOL) v. 6 web application (https://itol.embl.de; Letunic & Bork, 2024).

### Whole-genome bisulfite sequencing

For WGBS, *D. vitifoliae* ‘G0’ adults and ‘G0’ and ‘G1’ eggs were collected from leaf galls at 15-23 DPI and kept at -80 °C until DNA extraction. For ‘G1’, DNA was isolated from pooled eggs collected from ‘G1’ galls, i.e. progeny of the ‘generation 1’ adults used for RNA-seq. Galls from one plant were pooled together to make one biological replicate. DNA was extracted from 200-250 eggs or 15-20 adults per replicate using a CTAB protocol based on the method described by Mugford et al. (2020). The quality and quantity of DNA was tested by running it on a 2% agarose gel and measurement with a Qubit Fluorometer. We prepared 150 bp paired-end WGBS libraries and used Illumina NovaSeq at the UC Davis Genome Center (https://dnatech.ucdavis.edu/#C2). Library prep and sequencing for the ‘G0’ eggs and adults was performed similarly by Novogene using Illumina NovaSeq (https://www.novogene.com/us-en/services/research-services/epigenome-sequencing/whole-genome-bisulfite-sequencing/). On average, we generated 47.2 (± 1.20 s.d.) million paired-end reads per sample (Table S1). Sequencing data were uploaded to the NCBI Sequence Read Archive (SRA) repository under Bioproject accession PRJNA1211173.

### DNA methylation analysis

Read and base quality statistics were generated for raw and processed WGBS reads using fastqc (version 0.11.9) (https://github.com/s-andrews/FastQC). Raw reads were initially trimmed and filtered using Trim Galore (version 0.6.7) (https://github.com/FelixKrueger/TrimGalore) with default parameters plus an 80 bp minimal read length filter. Fastqc results showed the 5’ and 3’ ends of reads had artifactual base sequences that were not removed by quality trimming, and libraries had poor alignment efficiency when mapped to the bisulfite-converted reference genome (Rispe et al., 2020). Thus, 25 bp at the 5’ ends and 20 bp at the 3’ ends were trimmed from all reads using Trim Galore. Reads were mapped to the reference genome using bismark v. 0.24.1 (Krueger & Andrews, 2011). On average, 59.4 % (± 0.71 s.d) of reads uniquely mapped to the bisulfite-converted reference genome (Table S1). After removing PCR duplicate reads, the average coverage was 5.21 Gigabases (± 1.64 s.d), or 17.7x (± 5.59 s.d) genome coverage (Table S1). Genome-wide CpG, CHG, and CHH methylation percentages for each sample were extracted from the bismark alignment report and plotted using ggplot2.

To determine which CpG sites were significantly methylated, a binomial test was performed using the bisulfite conversion (C to T) error rate, which was 0.3% (± 0 s.d, n=4) based on the CpG methylation percentage of spiked-in lambda bacteriophage DNA. CpG sites were filtered based on a BH-adjusted p-value cutoff of 0.001, which removed CpG sites that had less than ∼10% methylation (methylated reads divided by all aligned reads). CpG sites were annotated with gene information (gene ID, functional group) and genome features (UTRs, introns, exons, intergenic) using the ‘intersect’ command of bedtools v. 2.30.0 (Quinlan & Hall, 2010). To assess global differences in CpG methylation between samples, principal component analysis (PCA) was performed using prcomp (R stats package v. 4.3.1). For PCA, we included only CpG sites that were covered by ≥4 reads in all samples, were methylated in at least one sample (binomial test, BH-adjusted p-value cutoff of 0.001), and had non-zero variance. To determine if any *D. vitifoliae* genes were differentially methylated after feeding on different hosts, we used a logistic regression with a F-test and applied a q-value cutoff of 0.05 and a ≥10% methylation difference cutoff using the R package methylKit v. 1.28.0 (Akalin et al., 2012). The ‘intersect’ command of bedtools v. 2.30.0 was used to determine if any differentially methylated genes were also differentially expressed (i.e. host-responsive).

Additionally, we wanted to understand how gene expression plasticity correlated with DNA methylation. We used all host-by-host contrasts to calculate the mean absolute value of log2-fold-change in gene expression as a measure of transcriptional plasticity. The ‘G0’ (starting colony) adult samples were used to calculate DNA methylation levels for each gene, since these were sampled at the same developmental stage (adults) as our RNA-seq data. Transcriptional plasticity and DNA methylation percentage (% reads with methylated cytosines divided by total aligned reads) was plotted for each gene in a scatterplot and a LOESS (Locally Estimated Scatterplot Smoothing) line was fitted to the data to visualize the trend. We then tested for correlation between transcriptional plasticity and DNA methylation with a Spearman rank correlation test using the R stats package v. 4.3.1. Lastly, genes were binned into quartiles based on the mean absolute value of log2-fold-change, then pairwise Wilcoxon rank-sum tests were performed to test for significant differences in DNA methylation levels between quartiles.

## Results

### Insect fitness differs significantly across host grapevine genotypes

To determine how fitness differs across grapevine hosts, we experimentally introduced a clonal line of *D. vitifoliae* to a panel of grapevine genotypes, including a non-native host (Cab33) and interspecific hybrids of native North American grapevine species (Figure 1). Fitness varied across host grapevine genotypes (Figure 1A) but the grapevine genotypes SW (*V. riparia* X *V. rupestris*), 5BB (*V. riparia* X *V. berlandieri*), and 3309 (*V. riparia* X *V. rupestris*) were incompatible, i.e. fully resistant to colonization. Attempted feeding by *D. vitifoliae* induced small necrotic lesions on leaves of genotypes SW and 5BB within six days after experimental infestation (Figure 1D and 1E). Interestingly, genotype 3309 showed symptomless resistance after six days (Figure 1F) but eventually showed necrotic lesions in leaves at feeding sites three weeks after infestation (Figure S1). Fitness was lower on both Cab33 and SO4 (*V. berlandieri x V. riparia*) compared to the other three susceptible hosts (Figure 1G). All other pairwise comparisons between host genotypes revealed no differences in *D. vitifoliae* fitness.

**Figure 1:**
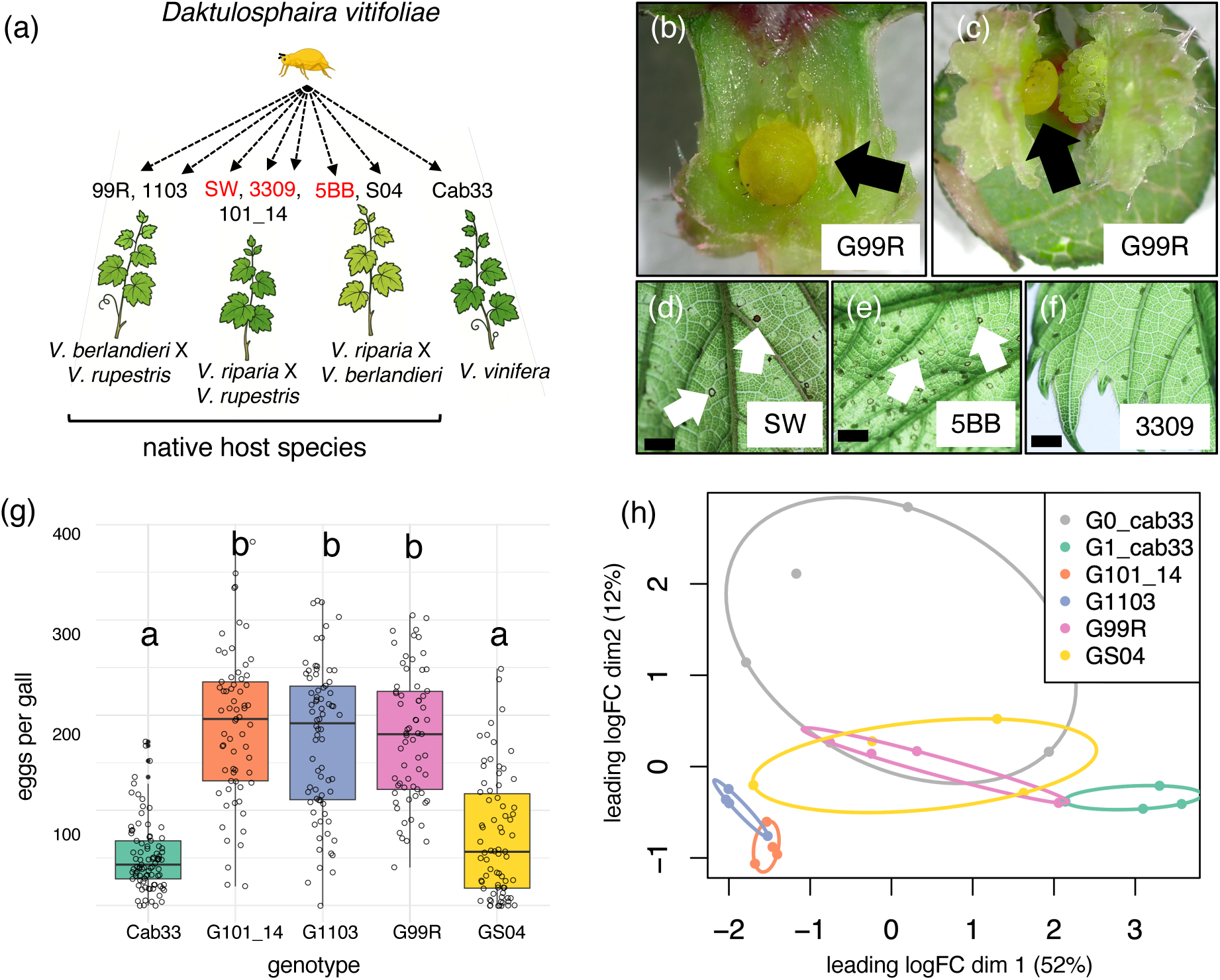
(a) Overview of the initial experimental setup; genotype names in red were resistant to *D. vitifoliae,* insects died and were excluded from downstream analysis. (b-c) *D. vitifoliae* adult (black arrows) within its gall at 3 weeks after infestation of host genotype 99R. (d-f) Young leaves of resistant host plants at 6 days after infestation; white arrows indicate sites of leaf necrosis induced by *D. vitifoliae* feeding/probing; horizontal black scale bars are 1 mm in length. (g) Box plot showing interquartile range and median (horizontal line within box) of *D. vitifoliae* fitness (eggs per gall) across susceptible host genotypes; genotypes with letters ‘a’ are significantly different from those with letter ‘b’ based on pair-wise Wilcoxon tests (Benjamini-Hochberg [BH]-adjusted p-values are all less than 0.0001). (h) Multi-dimensional scaling plot of log-CPM (gene counts per million gene counts) for all RNA-seq samples.

### *D. vitifoliae* gene expression is highly dynamic and responsive to host grapevine genotype

To understand the genome-wide variation in *D. vitifoliae* gene expression induced upon transfer to different host grapevine genotypes, we isolated RNA from *D. vitifoliae* adults after feeding on new hosts for a single generation and performed RNA-seq. Clustering of samples based on multidimensional scaling (MDS) indicated host environment induces distinct changes in *D. vitifoliae* gene expression (Figure 1H). Differential gene expression analysis revealed 3,706 were host-responsive (1.5-fold-change in any pairwise comparison, BH-adjusted p < 0.05; Table 1), which was 28% of the 13,125 total expressed genes. We then performed a Fisher’s exact test to identify gene functional groups that are enriched (over-represented) among the host-responsive genes relative to the genome-wide background, which included all 25,808 *D. vitifoliae* gene models. Genes encoding predicted secretory effectors (adj. p = 1.7e-127, odds ratio = 3.30), developmental regulators (adj. p = 1.6e-14, odds ratio = 4.55), cuticle- and chitin-associated proteins (adj. p = 1.5e-06, odds ratio = 3.03), detoxification enzymes (adj. p = 1.9e-05, odds ratio = 2.14), and proteins involved in general metabolism (adj. p = 8.9e-03, odds ratio = 5.23) were enriched among host-responsive genes (Table 1).

**Table 1:**
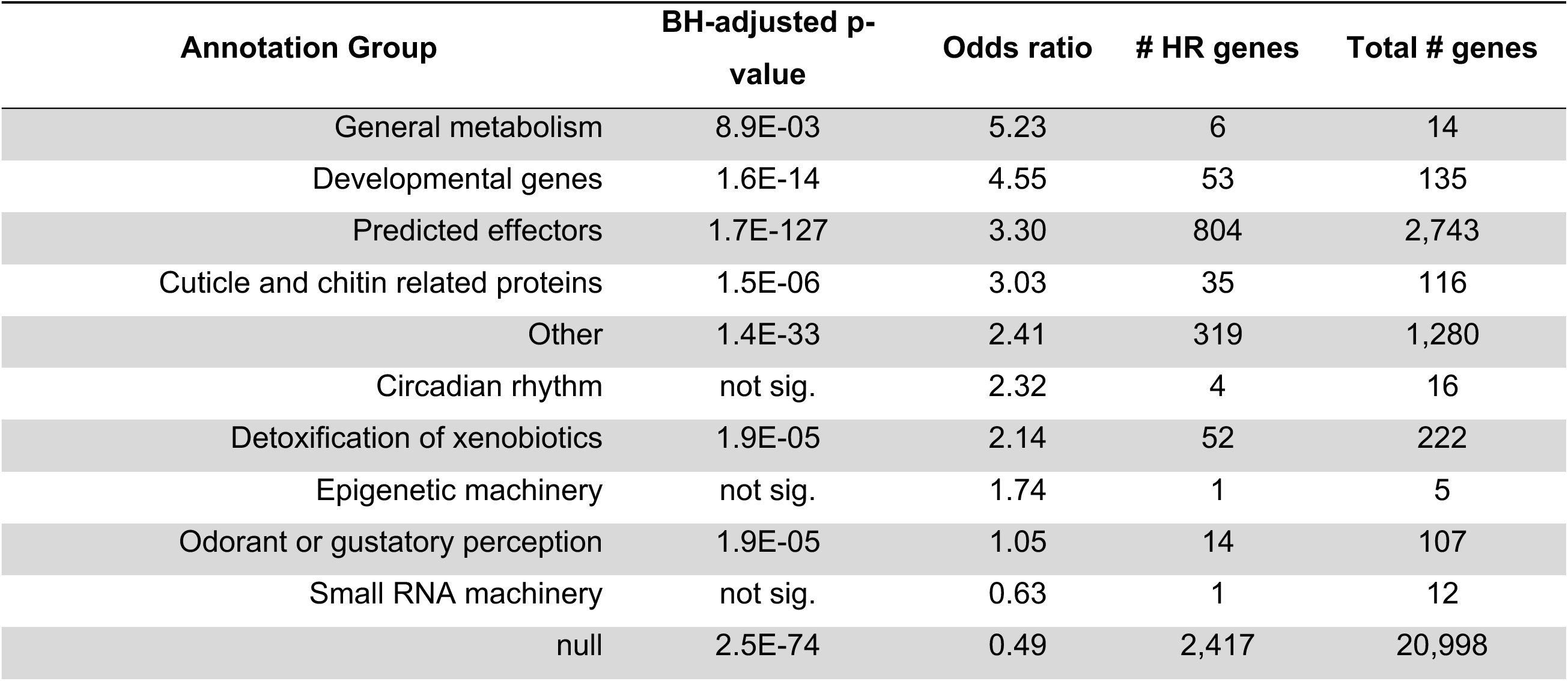
Fisher’s exact test for functional group enrichment among host-responsive (HR) genes.

Pair-wise analysis of gene differential expression (DE) revealed that the non-native host Cab33 induced the strongest change in gene expression compared to other hosts; 3,432 (92.6%) of the total 3,706 host-responsive genes show differential expression in contrasts between Cab33 and the other four host genotypes (Figure 2A). Only 19 genes were differentially expressed in all contrasts with Cab33 (Figure 2B); 6 of these genes encoded predicted secretory effectors, including 3 RING-type E3 ligases (DV3020556, DV3022321, and DV3025927). We further investigated the types of genes that were up- and down-regulated when *D. vitifoliae* feeds on Cab33 using gene enrichment analysis. Of the host-responsive genes, 2,667 (72%) were upregulated during interaction with Cab33 relative to all other hosts. Genes encoding predicted effectors, cuticle-and chitin-associated proteins, and detoxification enzymes were enriched among genes upregulated on Cab33 (Table S2). Enriched KEGG terms for upregulated genes included glutamate receptors, fatty acyl-CoA reductases, cathepsin B cysteine proteases, nicotinic acetylcholine receptors, and organic acid transporters (Figure 2C). Genes downregulated during interaction with Cab33 are enriched for functions related to development (Fisher’s exact test, BH-adjusted p = 1.59e-09, odds ratio = 6.36) and KEGG terms for lactose galactohydrolase/β-glucosidases (PFAM domain GH1) and serine protease inhibitors (Figure 2D). Overall, these results suggest that the *D. vitifoliae* transcriptome is highly plastic, especially at genes encoding secretory proteins including putative effectors and genes involved with response to and protection against environmental stress.

**Figure 2:**
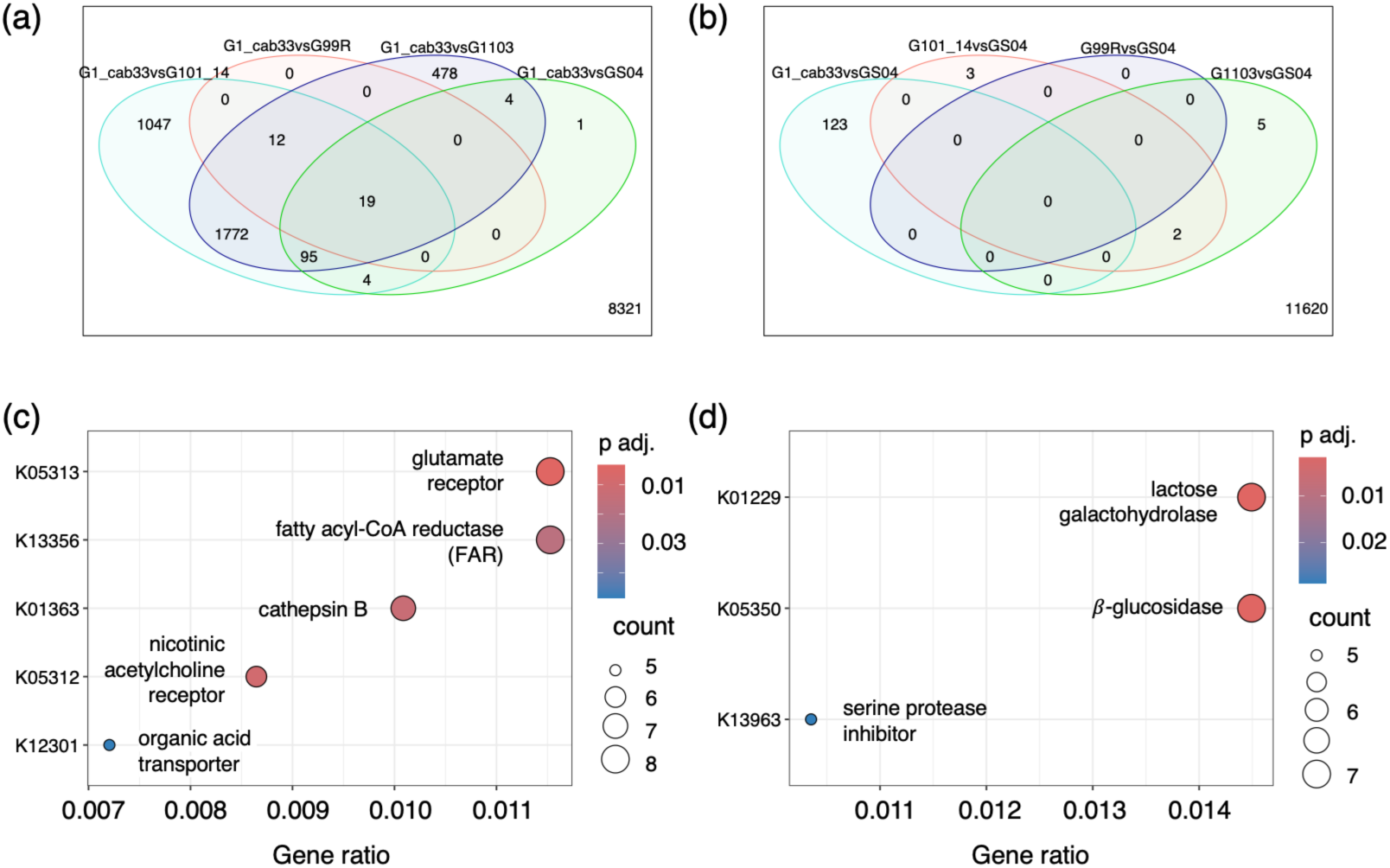
Transcriptional plasticity induced by host transfer. (a-b) Venn diagrams showing the number of differentially expressed *D. vitifoliae* genes contrasting *V. vinifera* cv Cab33 (a) or SO4 (b) with all other hosts genotypes. (c-d) KEGG terms enriched among genes upregulated (c) and downregulated (d) during interaction with *V. vinifera* cv Cab33 compared to all other hosts; colors represent Benjamini-Hochberg (BH) adjusted p-values.

### Four gene co-expression modules co-vary with *D. vitifoliae* fitness

To identify groups of genes that likely function together in shared biological processes, we performed weighted gene co-expression network analysis using 13,917 genes after removing ‘zero genes’ that showed negligible expression. A signed gene co-expression network was constructed that contained 88 modules based on hierarchical clustering of gene connectivity dissimilarity (Figure 3A and 3B). Gene co-expression modules were named based on colors randomly assigned by the WGCNA program (Langfelder & Horvath, 2008). To identify groups of co-regulated genes (modules) that might underlie differences in *D. vitifoliae* fitness across host genotypes, we tested for correlation between module eigengenes (first principal component of the module and the most representative gene) and fitness using linear regression tests. Five eigengenes significantly correlated with *D. vitifoliae* fitness (Figure 3C-F); we excluded the ‘grey’ module from further analysis since it is a bin for genes that cannot be assigned to co-expression modules and are unlikely to be co-regulated.

**Figure 3:**
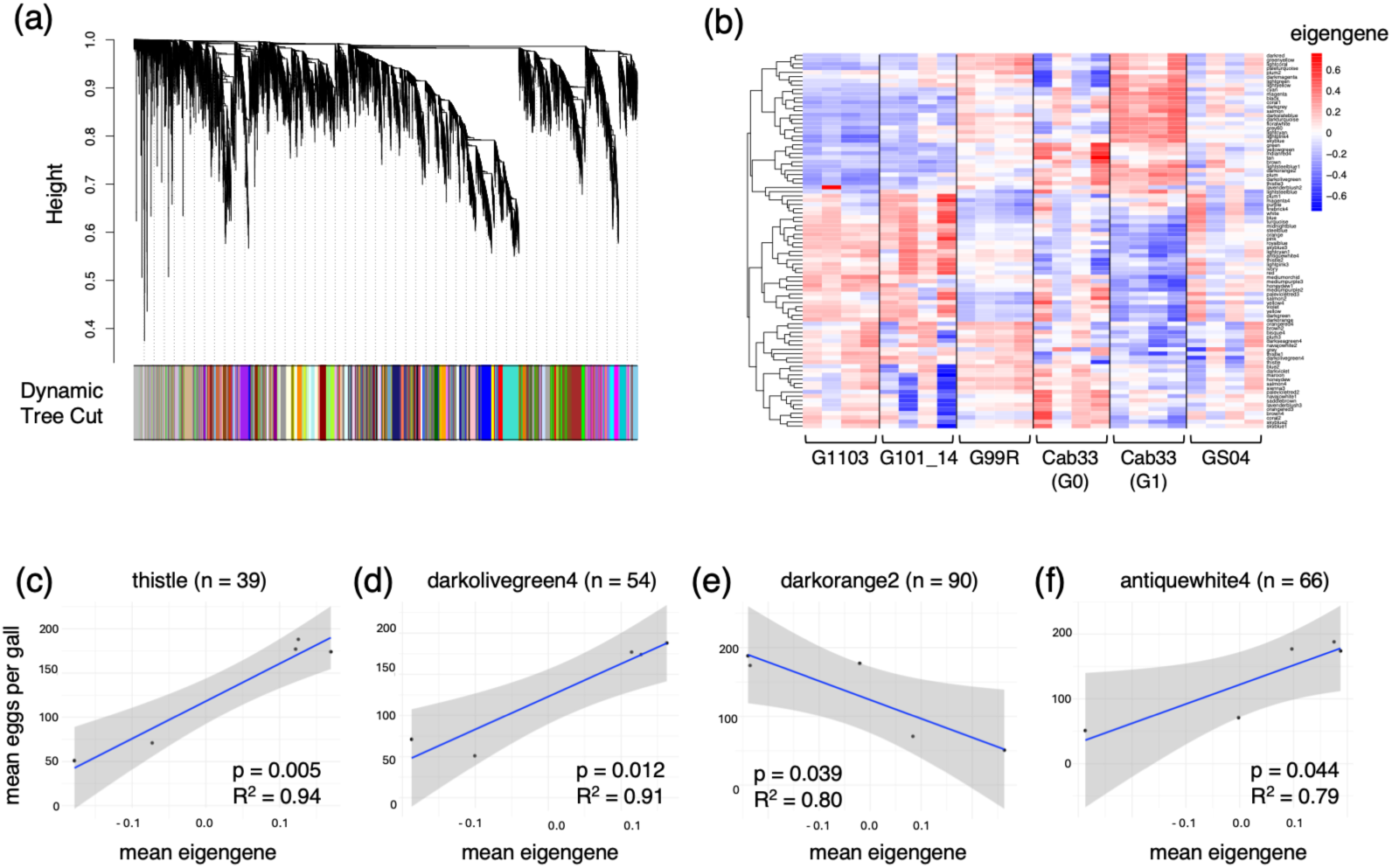
Gene co-expression network analysis. (a) Gene dendrogram for *D. vitifoliae* signed (unidirectional) gene co-expression network and module assignment (bottom). (b) Heatmap of module eigengenes (first principal component of the module, the ‘most representative’ gene) for all 88 modules (vertical axis) clustered based on Euclidean distance. (c-f) Scatter plots and results of linear regression tests for four co-expression modules that correlate in expression with *D. vitifoliae* fitness.

We performed Fisher’s exact tests to identify gene co-expression modules that were enriched for host-responsive genes. Twenty-one out of the 88 modules were enriched for host-responsive genes (odd ratio >2, BH-adjusted p <0.05) (Table S3), which reflects the high number of transcriptionally plastic genes in the dataset. The ‘darkorange2’ module was both enriched for host-responsive genes and negatively correlated with *D. vitifoliae* fitness (Figure 3E). Based on KEGG term enrichment analysis, ‘darkorange2’ was overrepresented for genes that encode glycoside hydrolase family 1 enzymes (PFAM GH1), and showed distinct patterns of expression across host genotypes (Figure S3). A small cluster of 24 genes was highly expressed during interactions with native hosts 101-14 and 1103 but decreased in expression during interactions with Cab33 and SO4, which are hosts that maintain relatively low *D. vitifoliae* fitness. The remaining 66 genes in the module are grouped into a larger cluster that shows the opposite trend in expression, largely increasing in expression during interaction with Cab33 and SO4 relative to 101-14 and 1103. These results show the ‘darkorange2’ module is highly responsive to host environment and may contribute to fitness variation in *D. vitifoliae* during interaction with non-optimal hosts.

### One module contains a cluster of conserved and proline-rich secretory proteins

The ‘thistle’ module was most strongly correlated with *D. vitifoliae* fitness (R^2^ = 0.94, p = 0.005; Figure 3C-F) and enriched with genes encoding predicted secretory effectors (Fisher’s exact test, BH-adjusted p = 0.028, odds ratio = 3.32). Other than the 11 predicted effectors, the remaining 28 genes in this module did not belong to any predicted functional group. Four predicted effectors were homologous to *Acyrthosiphon pisum* ‘lysine-rich arabinogalactan protein 19-like’ (accession XP_001948476.1), based on previously published BLASTp alignments (Rispe et al., 2020). We termed these genes as encoding ‘<u>AR</u>abino<u>G</u>alactan-<u>L</u>ike’ (ARGL) proteins. *ARGL* genes show similar patterns of transcript expression across treatments, generally increasing in expression during interaction with high-fitness native hosts 101-14, 1103, and 99R relative to Cab33 (Figure 4B). We also noticed that the *ARGL* genes are located in a tandem cluster in the genome on scaffold 40 (Figure 4A). Within this same ∼50-kb spatial cluster are two more predicted effector-encoding genes (DV3003881 and DV3003883) and three ‘null’ genes (DV3003873, DV3003874, DV3003879); two of the three null genes (DV3003873 and DV3003874) are also assigned to the ‘thistle’ module, suggesting they are co-regulated with *ARGL* genes. To determine if *ARGL* genes have highly similar sequences and shared protein motifs, we performed Multiple Sequence Alignment (MSA) of the seven ARGL proteins using Clustal Omega (Sievers & Higgins, 2018). A sequence logo plot of the ARGL protein MSA showed that the proteins have a high proportion of proline residues and are highly conserved, especially at proline residues (Figure S4). We then constructed phylogenetic trees from the MSA of protein sequences and coding sequences (CDS) for all 2,743 predicted secretory effector proteins and found *ARGL* coding sequences clustered together into a monophyletic group (Figure S5), suggesting they belong to a gene family that evolved through tandem duplication. Despite being highly conserved at the amino acid level and clustered together based on coding sequences, ARGL protein sequences did not cluster together in a monophyletic group (Figure S6). Thus, ARGL proteins may have some structural and functional divergence from one another.

**Figure 4:**
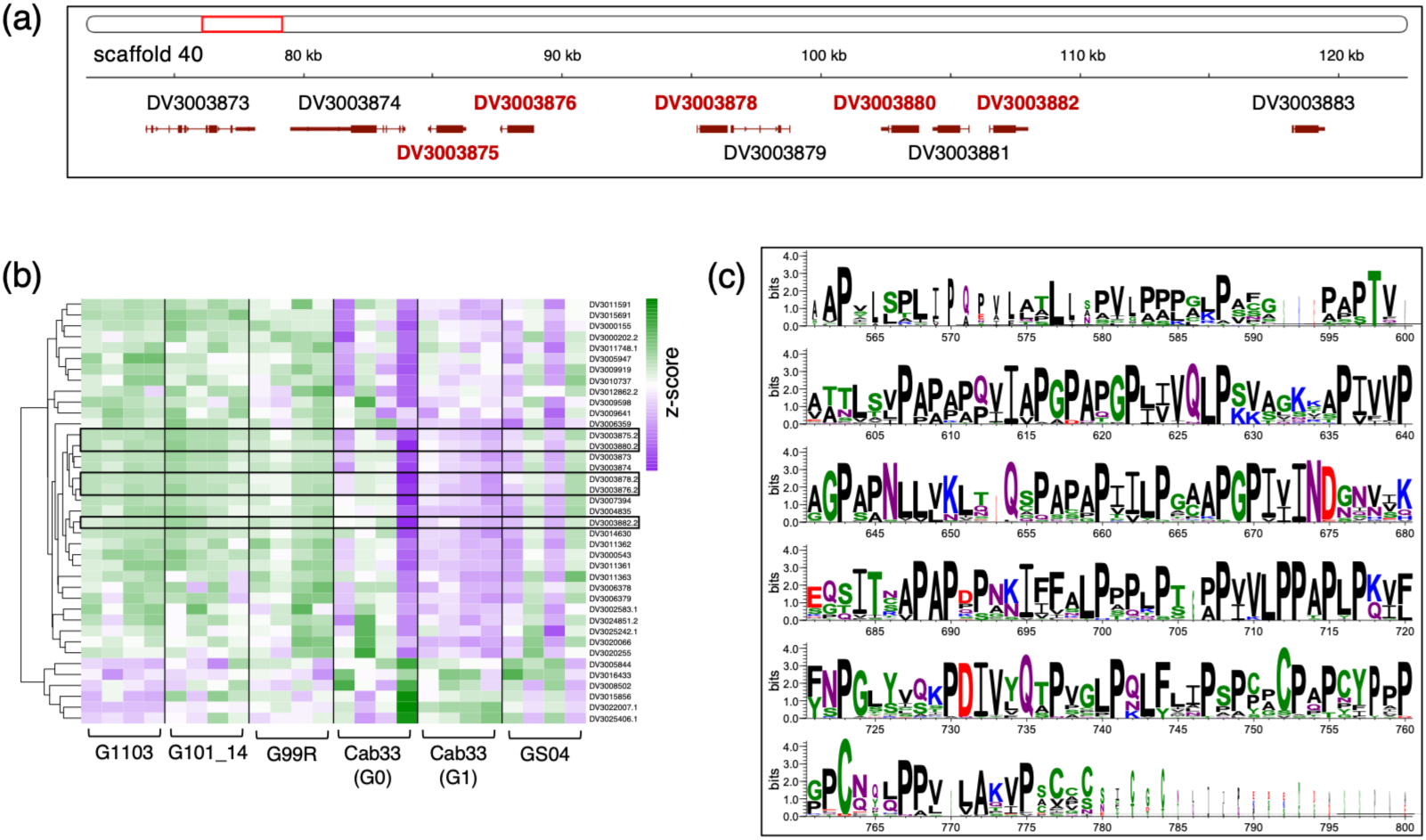
Spatial organization and gene expression profile of ARGL genes and multiple sequence alignment of ARGL proteins. (a) Genome browser view showing the ARGL locus in the *D. vitifoliae* genome; ARGL genes assigned to the thistle gene co-expression module are highlighted with bold red text. (b) Heatmap of normalized gene expression for genes in the thistle module; black rectangles highlight the ARGL genes. (c) Sequence logo plot based on multiple sequence alignment (MSA) of the seven predicted effector proteins encoded at the ARGL locus and the top seventeen orthologs of ARGL protein DV3003875; letters represent amino acid residues and letter size denotes the degree of conservation (larger letters are more conserved).

To identify putative orthologs of *ARGL* genes, we performed a BLASTp search using the *ARGL* gene DV3003875 queried against the NCBI non-redundant protein sequences (nr) database (https://blast.ncbi.nlm.nih.gov/Blast.cgi?PAGE=Proteins). We then performed reciprocal BLASTp alignments of the top 20 BLASTp hits against the *D. vitifoliae* genome. Seventeen out of the 20 proteins produced reciprocal best hits for the DV3003875 *ARGL* gene. We then investigated the evolution of the *ARGL* locus by performing MSA of the seven ARGL proteins with the 17 putative orthologs of ARGL DV3003875, which spanned adelgids (*Adelges cooleyi*) and eight aphid species (Table S4). Remarkably, a roughly 200-amino-acid region of all 27 proteins is highly conserved, especially at proline residues (Figure 4C and Figure S7). We also searched the proteins for predicted functional domains using InterPro (https://www.ebi.ac.uk/interpro/search/sequence/). Notably, all 24 proteins contain predicted secretory signal peptides but generally lack any known functional domains, with two exceptions. The *Aphis glycines* ‘hypothetical protein AGLY_000967’ (accession KAE9545424.1) and the *Metopolophium dirhodum* ‘uncharacterized protein LOC132951494’ (accession XP_060879307.1) contain the InterPro-unintegrated motif (PRINT entry) PR01217 (Proline-rich extensin signature). Extensin proteins are notably characterized as cell wall-associated proteins in plants (Moussu & Ingram, 2023), although whether the PR01217 motif confers any similar function to plant extensin proteins is unclear. Overall, our analysis of the ‘thistle’ co-expression module revealed an under-described group of genes encoding predicted secretory proteins, that are tightly co-regulated and positively correlated with fitness at the transcript level. ARGL proteins also are proline-rich, highly conserved, and belong to a putative family of secretory proteins seemingly widespread in the Aphidomorpha infraorder of Hemipteran insects. Together, these results suggest *ARGL* genes play an important role in host use in aphidomorph insects.

### DNA methylation in *D. vitifoliae* is mostly stable

In addition to our analysis of the *D. vitifoliae* transcriptome, we analyzed genome-wide patterns of DNA methylation using WGBS. Our goal was to first understand the general pattern of DNA methylation in the *D. vitifoliae* epigenome and then to determine whether host genotype induced changes in DNA methylation after insects fed on different hosts for a single generation. DNA methylation in *D. vitifoliae* occurs exclusively in the CpG context, and about 3.5-4% of CpG sites are methylated (Figure 5A), which is similar to what is reported in other Hemipteran insects (Bewick et al., 2016; Mathers et al., 2019). CHH and CHG methylation rates were around 0.3%, which is equal to the false positive rate. As in other insects, DNA methylation in *D. vitifoliae* occurs mostly in exons and other gene body regions (Figure 5B).

**Figure 5:**
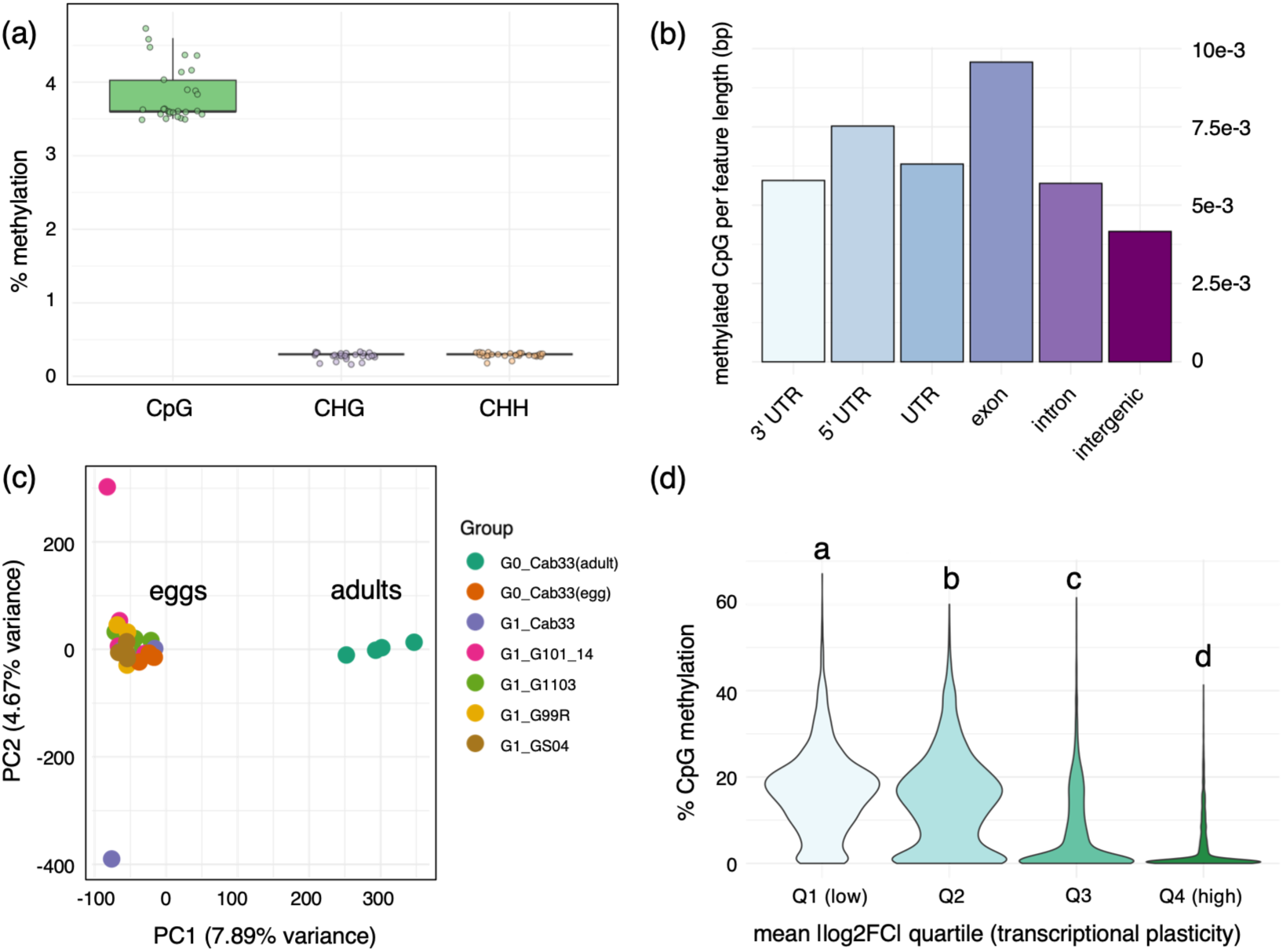
DNA methylation in *D. vitifoliae*. (a) Box plot showing the proportion of methylated cytosines (vertical axis) for each sequence context (CpG, CHG, and CHH). (b) Bar plot showing CpG methylation levels in adults for each genomic region (untranslated regions (UTRs), intron, exon, and intergenic); the vertical axis is the total number of methylated CpG sites divided by the total length (in base pairs) of the genomic region. (c) Principal component analysis (PCA) based on methylation levels at 201,448 CpG sites significantly methylated in at least one sample and covered by 4 or more reads in all samples. (d) Width-scaled violin plots showing CpG methylation levels in adults (vertical axis) per transcriptional plasticity quartile; genes that were expressed and had CpG methylation data (n = 12,368) were binned into quartiles based on mean absolute value of log2-fold-change in expression (a quantification of transcriptional plasticity induced during transfer to different hosts); letters above the quartiles indicate significant difference based on pair-wise Wilcoxon tests (all p-values < 2e-16).

To determine variation in DNA methylation across our samples at a global level, principal component analysis (PCA) was performed using the 201,448 CpG sites that were methylated in at least one sample and had non-zero variance. Our PCA revealed no distinct clustering based on treatment group (host grapevine genotype) except for the adult samples, which were distinctly subclustered away from the remaining samples (n=24), all of which were collected at the egg stage. (Figure 5C). We repeated the PCA twice: once after removing the two potential outlier samples, and then once more after further removing the adult samples. The results similarly showed no distinct clustering based on host genotype (Figure S8). We additionally tested whether any genes were differentially methylated between insects reared on different host grapevine genotypes using methylKit (Akalin et al., 2012). Eight genes were host-responsive at the DNA methylation level (Table S5), but none of these genes were differentially expressed at the transcript level. All 3,706 genes that were transcriptionally plastic (host-responsive) showed no significant change in DNA methylation in the subsequent generation. These results show that host-induced variation in gene expression does not lead to any significant change in gene body methylation in progeny.

### Gene transcriptional plasticity is negatively correlated with DNA methylation

We wanted to further understand how host-induced transcriptional plasticity correlates with DNA methylation in *D. vitifoliae* adults. To quantify the degree of transcriptional plasticity for genes, mean absolute values of log2 fold changes in gene expression across all host-by-host contrasts were calculated. Gene-level transcriptional plasticity and DNA methylation level was plotted for genes that were expressed and had CpG methylation data in adults (n = 12,368) (Figure S9). There was a significant negative correlation between transcriptional plasticity and DNA methylation (Spearman’s *ρ* = -0.54, p < 2.2e-16). To further visualize the correlation between gene expression plasticity and DNA methylation, genes were binned into quartiles based on transcriptional plasticity and pair-wise Wilcoxon rank-sum tests were performed. This also showed a negative correlation between transcriptional plasticity and DNA methylation, with transcriptionally plastic genes having significantly less methylation compared to stably expressed genes (Figure 5D). These data indicate gene body DNA methylation contributes to gene expression regulation in *D. vitifoliae* and may stabilize gene expression levels when *D. vitifoliae* faces new host environments.

## Discussion

The transcriptional and epigenetic mechanisms that enable parasites and pathogens to colonize of new hosts remain only partially understood. In this study, we found the transcriptome of the specialist insect *D. vitifoliae* is remarkably plastic, but more so when colonizing a novel (non-native) host compared to native hosts. The transcriptionally plastic genes that increased in expression were enriched for predicted secretory effectors, detoxification enzymes, and cuticle- and chitin-associated proteins – genes linked to the herbivore lifestyle (Karban, 2020; Peláez et al., 2023). We found gene co-expression modules correlated with fitness, suggesting plasticity in transcriptional networks contributes to *D. vitifoliae* adaptation to host plants. Our analysis of DNA methylation in adults revealed a negative correlation between gene-level CpG methylation and host-induced transcriptional plasticity. These results highlight that rapid transcriptional change may enable host shifts in a specialist herbivore similar to how polyphagous herbivores adapt to feed on divergent hosts (Dermauw et al., 2013; Mathers et al., 2017; Snoeck et al., 2018). These results also link a lack of DNA methylation to transcriptionally plastic genes and suggest gene body methylation plays an important role in constitutive gene expression during colonization of new hosts. Together these results help us understand how transcriptional plasticity and gene body DNA methylation enable acclimatization to novel hosts by specialist herbivores.

Because *D. vitifoliae* is a specialist herbivore that evolved to feed on a narrow range of hosts in the *Vitis* genus, we predicted that transcriptional plasticity induced by transfer to different compatible hosts would be low; this has been shown for other specialist insect species in similar experiments (Govind et al., 2010; Q. Yu et al., 2016; Roy et al., 2016; Birnbaum et al., 2017; Schweizer et al., 2017). Contrary to this prediction, the *D. vitifoliae* transcriptome was highly plastic. Of all expressed genes, 3,706 (28%) were host responsive (Table 1), and the majority of these were upregulated while feeding on the non-native host European wine grape *V. vinifera* (cv Cab33) (Table S2). Predicted detoxification genes were enriched among genes upregulated during interaction with *V. vinifera*, and included numerous cytochrome P450s, glutathione S-transferases and UDP-glycosyltransferases, and ABC transporters. These patterns suggest that all three phases of xenobiotic detoxification are reprogrammed when *D. vitifoliae* feeds on *V. vinifera* (Heidel-Fischer & Vogel, 2015), likely helping to overcome novel host defenses. Predicted effectors were also enriched among genes upregulated while feeding on *V. vinifera* (Table S2). This functional group includes over 2,700 secretory proteins with effector attributes, including putative effectors known to interact with host grapevine proteins such as RING-type E3 ligases (Zhao et al., 2019). Interestingly, three RING-type E3 ligases were consistently upregulated when *D. vitifoliae* fed on the non-native host *V. vinifera* compared to the other hosts, supporting the hypothesis that protein modification is a key pathway to coopt hosts (Zhao et al., 2015; Huang et al., 2021). The subset of genes identified here serve as targets for further functional characterization to validate mechanisms of host adaptation in *D. vitifoliae* and improve management of grape against *D. vitifoliae.* More broadly, these results show that at least some lineages of specialist herbivores, despite long-term ecological specialization, rely on substantial transcriptional plasticity for ecological persistence, contrary to predictions (Sexton et al., 2017; Birnbaum & Abbot, 2020).

Our analysis of KEGG terms similarly revealed that genes putatively involved with insect adaptations were overrepresented among genes upregulated while feeding on the non-native host *V. vinifera*. These genes included ionotropic glutamate receptors which can function as chemosensory receptors (Benton et al., 2009; Croset et al., 2010), fatty acyl-CoA reductases (FAR) which can modify cuticle composition (Li et al., 2020; Di et al., 2023), and cathepsin B cysteine proteases (Figure 2C). Notably, aphid cathepsin B proteases function as secretory effectors during interaction with divergent host plants (Mathers et al., 2017). Other recent studies show *M. persicae* cathepsin B genes are expressed in response to abiotic stresses and important for thermotolerance (Liu et al., 2023; Zhang et al., 2025). Thus, cathepsin B proteins in related aphids may contribute to adaptation to different environments and may also underlie host adaptation in *D. vitifoliae*. In summary, we found many of the genes that increase in expression during colonization of a non-native host are putatively involved with key adaptations in herbivorous insects – detoxification of plant defense metabolites, direct manipulation of hosts via effector protein secretion, and cuticle modification – highlighting that specialist lineages dramatically reprogram critical pathways to colonize new environments.

In addition to differential expression analysis, we wanted to understand how transcriptional plasticity mediates host use by correlating clusters of co-expressed genes (gene co-expression modules) with variation in *D. vitifoliae* fitness across different hosts. Four gene co-expression modules were significantly correlated with variation in fitness, indicating that plasticity in transcriptional networks may contribute to host plant adaptation (Figure 3). The ‘thistle’ co-expression module most strongly correlated with fitness (Figure 3C) and was enriched with genes coding for predicted secretory effector proteins (Fisher’s exact test, BH-adjusted p = 0.028), while the remaining genes in the module were not assigned to any predicted functional group. This module also contained several co-regulated genes that are organized as a spatial cluster in the genome, encode predicted secretory effector proteins (ARGL; Figure 4), and are shared among aphids indicating their likely presence in the common ancestor of the Aphidomorpha lineage. *D. vitifoliae* ARGL proteins and putative ARGL orthologs show a high degree of amino acid conservation and retained secretory function over the course of more than 160 million years of evolution (Figure 4C and Table S4)(Ren et al., 2013), highlighting an important role for ARGL proteins in Aphidomorph biology or ecology. Some of the putative ARGL orthologs in aphids and adelgids (*Adelges cooleyi*) contain predicted proline-rich extensin motifs, and other putative ARGL orthologs are annotated as ‘lysine-rich arabinogalactan protein 19-like’, ‘vegetative cell wall protein gp1-like’, and ‘leucine-rich repeat extensin-like protein 1’ (Table S4). Although the function of insect ARGL proteins remains unclear, a common theme among annotations is the sensing and maintenance of plant cell walls (Herger et al., 2019; Ellis et al., 2010; Ferris et al., 2001). As a central regulator of cellular structure and immune signaling in plants, the plant cell wall is a critical interface for herbivore-host interaction (Wolf, 2022; Erb & Reymond, 2019), and thus a likely target of insect secretory effectors. Whether ARGL proteins play a role in host cell wall manipulation or host interaction more broadly requires further functional genetic analysis, although our computational analyses here suggest they are important for host use across aphidomorph insects.

In addition to our analysis of transcriptomic plasticity induced by host transfer, we analyzed DNA methylation in *D. vitifoliae* adults (initial generation, G0) and in eggs after one generation on different host plant genotypes. To our knowledge, no other studies have measured plasticity in both the transcriptome and epigenome of an herbivore during interaction with novel hosts. Thus, we lack data for how host plants shape the epigenome of herbivores and on the correlation between gene expression variation and epigenetic marks. Importantly, our data were collected across two different generations of *D. vitifoliae*. We collected RNA-seq data from adult *D. vitifoliae* that fed on new host plants for one generation and DNA methylation in the progeny eggs. While this experimental design limits our ability to correlate individual-level gene expression with DNA methylation, our cross-generation comparison allowed us to test for correlation between host-induced transcriptional plasticity and variation in DNA methylation in the subsequent generation. Although the *D. vitifoliae* transcriptome was highly plastic across host plants, there was little variation in DNA methylation in the subsequent generation. We found eight genes that were differentially methylated across host plants (Table S5); thus, we did find evidence for host-specific epimutations that could affect host adaptation via change in gene expression level (Bonasio et al., 2012; Libbrecht et al., 2016; Mathers et al., 2019), transcript splicing (Li-Byarlay et al., 2013; Glastad et al., 2016), or gene mutation rate (Elango et al., 2009). However, none of these eight genes were differentially expressed across hosts in the parent (adult) generation or linked to fitness variation and samples did not cluster based on host genotype (Figure 5C and Figure S8). These results show that host-induced plasticity in gene expression does not correlate with changes in DNA methylation in the subsequent generation, suggesting changes in DNA methylation are unlikely to mediate rapid phenotypic adaptation. We found a significant negative correlation between transcriptional plasticity and DNA methylation in adults, however, with transcriptionally plastic genes being generally depleted in DNA methylation relative to stably expressed genes (Figure 5D and Figure S9). This is consistent with previous findings in other insects (Bonasio et al., 2012; Libbrecht et al., 2016; Mathers et al., 2019). Given the adaptive significance of transcriptional plasticity during host use and host shifts, DNA methylation may play a role by limiting plasticity of stably expressed genes for which environment-induced plasticity is maladaptive. Future experiments are still needed to validate the role of DNA methylation in mediating transcriptional plasticity during host plant acclimatization.

Taken together, our results suggest that specialist herbivores, despite long-term ecological specialization, can widely and rapidly reprogram their transcriptomes to feed on new hosts, and that gene body methylation contributes to the regulation of host-induced transcriptional plasticity. In this example, the switch to the novel host *V. vinifera* comes at a fitness cost but *D. vitifoliae* still persists, highlighting a mechanism for adaptation to increased host breadth that may occur with invasions and range expansions. More broadly, these data can serve as the basis for future functional genetic work to validate genes involved with adaptation to host plants in aphidomorph insects.

## Data availability

Sequencing data are publicly available on NCBI Sequence Read Archive (SRA) repository under Bioproject accession PRJNA1211173. All scripts for data processing and bioinformatic analyses are publicly available online at https://github.com/wcoll010/Grape-phylloxera-phenotypic-plasticity-during-host-transfer.

## Acknowledgments

We thank Conner Lay for help with collection of phenotypic data, Hung D. Lai for greenhouse maintenance, and Holly Clark, Clay Clark, and Wei Zhang at the UC Riverside genomics core for RNA library preparation. This project was supported by the following grants: USDA-NIFA Hatch grant CA-R-BPS-5199-H and CDFA grant CDFA-IAB #21-0426-000-SA awarded to PDN; The National Institute of General Medical Sciences of the National Institutes of Health grant no. R35GM151194 awarded to SCG.

## Author contributions

W.N.C., S.A.Z. and P.D.N. conceived the study. W.N.C. and S.A.Z. performed sample collections and insect performance assays. S.A.Z. performed the DNA and RNA extractions. W.N.C. performed data processing and analysis and wrote the manuscript with advice from S.C.G and P.D.N. All authors read, edited, and approved the manuscript.

---

**Figure S1:**
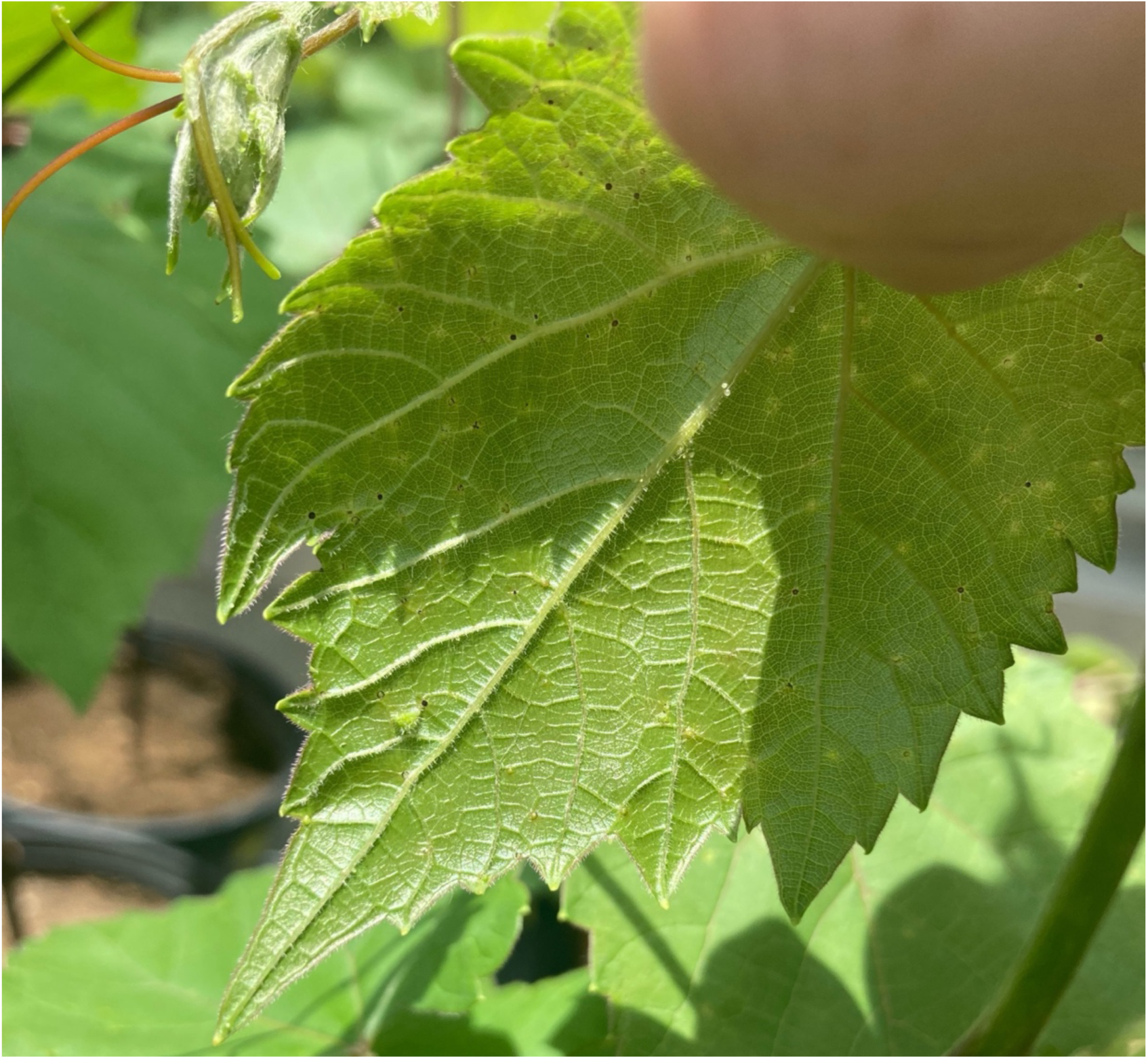
The underside of a 3309 (*V. riparia* x *V. rupestris*) leaf 3 weeks after infestation with *D. vitifoliae*. Small black necrotic lesions formed on the leaves where *D. vitifoliae* attempted to feed.

**Figure S2:**
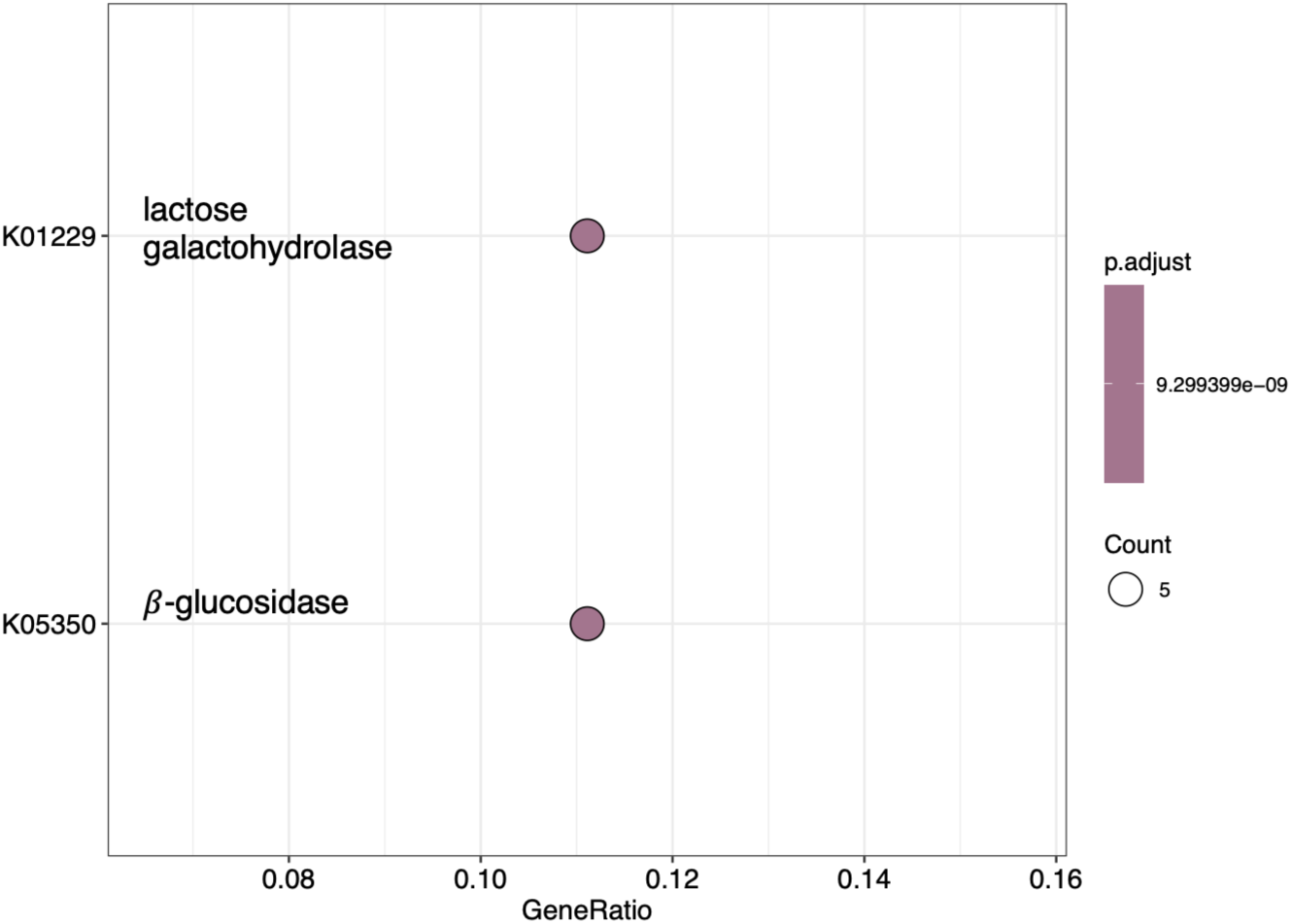
KEGG term enrichment analysis for genes in the darkorange2 co-expression module.

**Figure S3:**
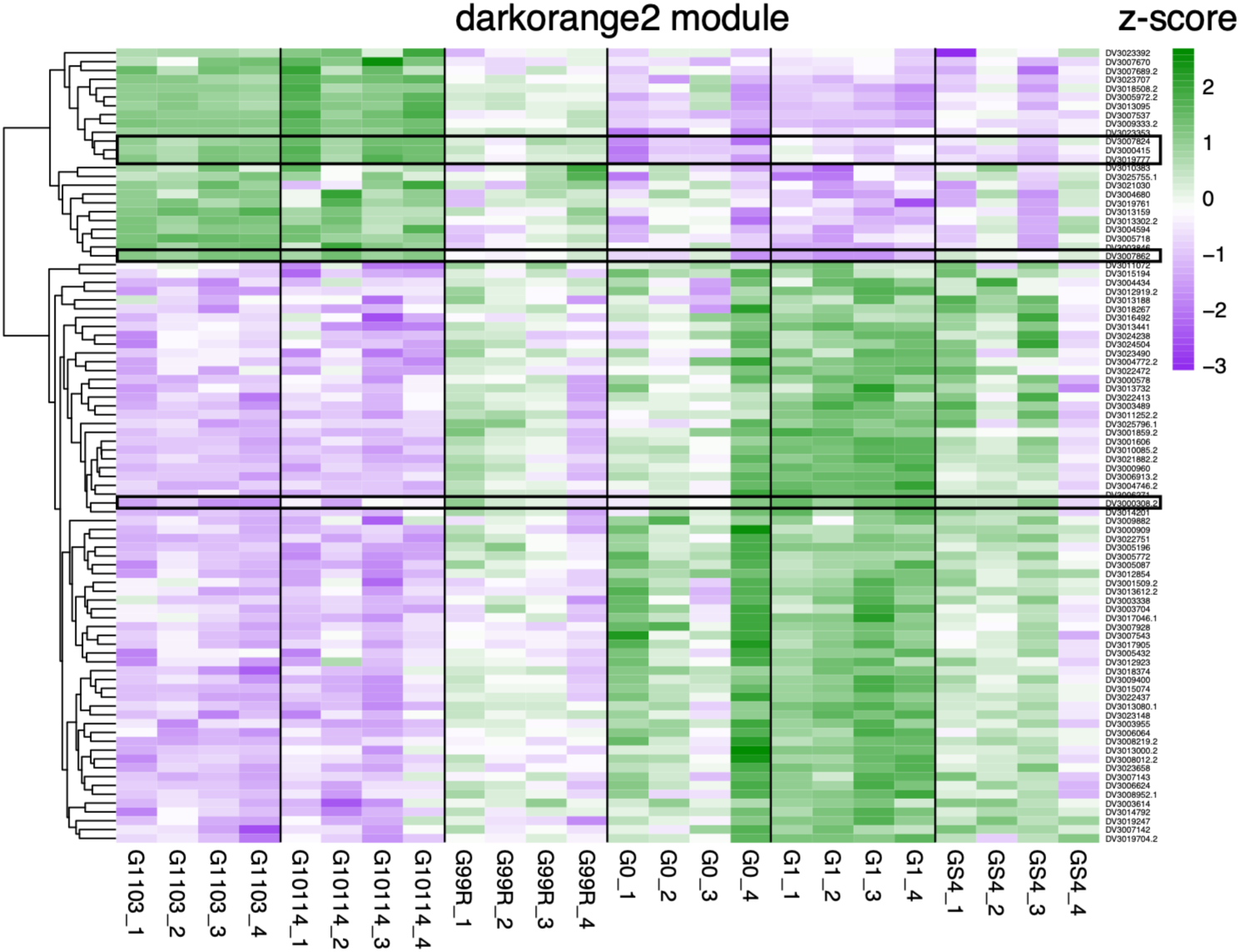
A heatmap of normalized gene expression for genes in the darkorange2 module. Black rectangles highlight genes with glycoside hydrolase KEGG annotation.

**Figure S4:**
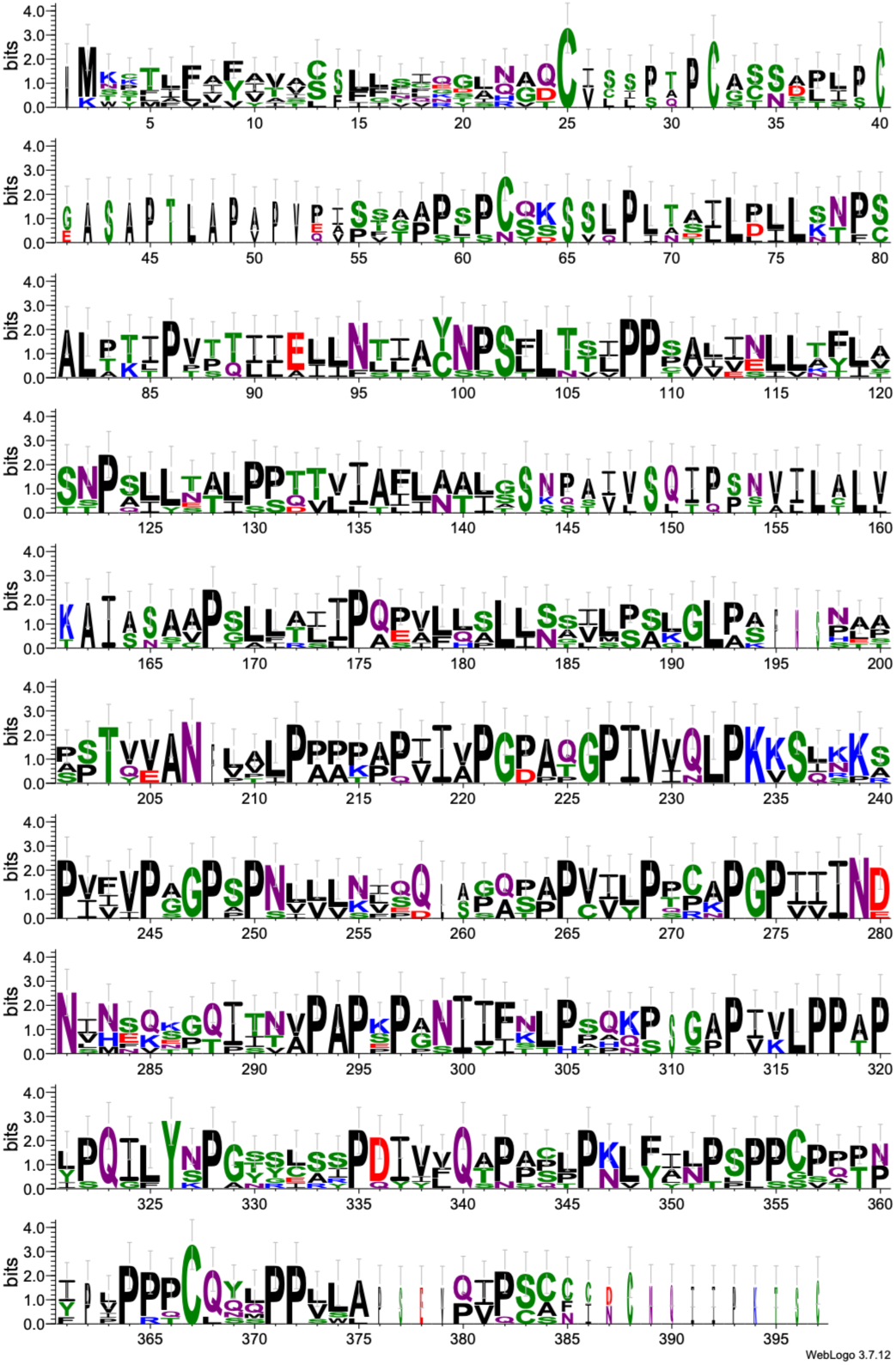
Sequence logo plot based on multiple sequence alignment (MSA) of the seven predicted effector proteins at the ARGL locus.

**Figure S5:**
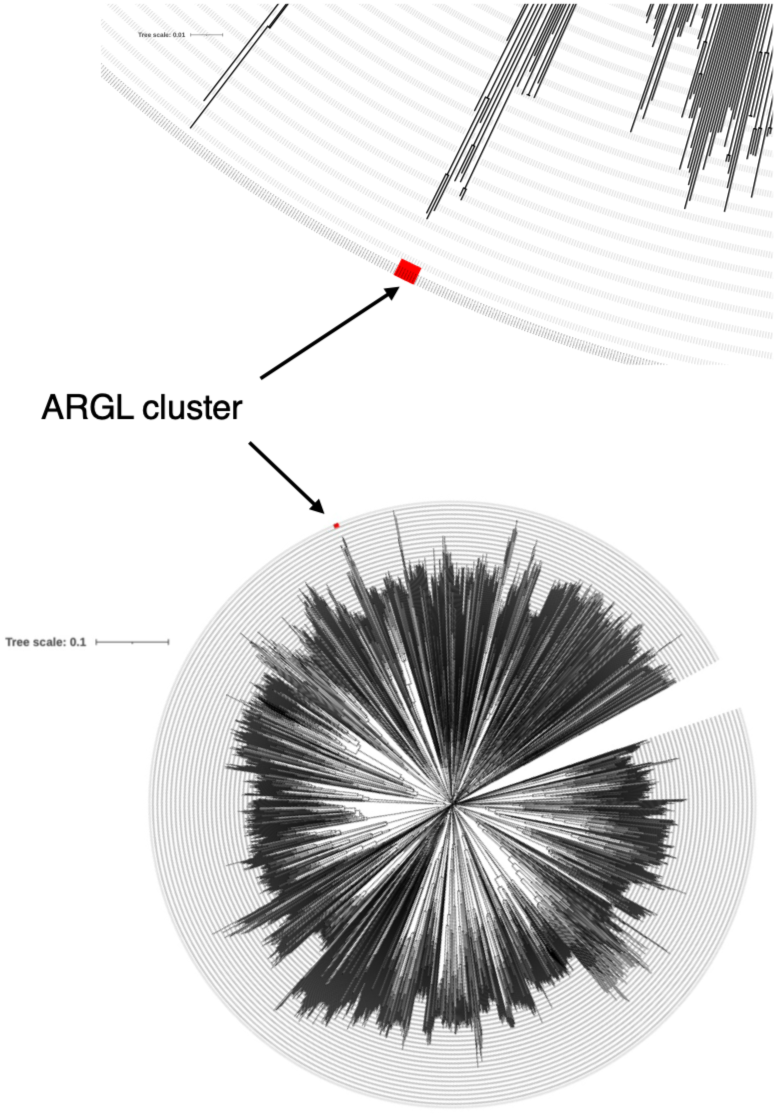
Phylogenetic tree of all 2,743 predicted *D. vitifoliae* effectors based on multiple sequence alignment (MSA) of coding sequences (CDS). ARGL coding sequences are highlighted in red with black arrows.

**Figure S6:**
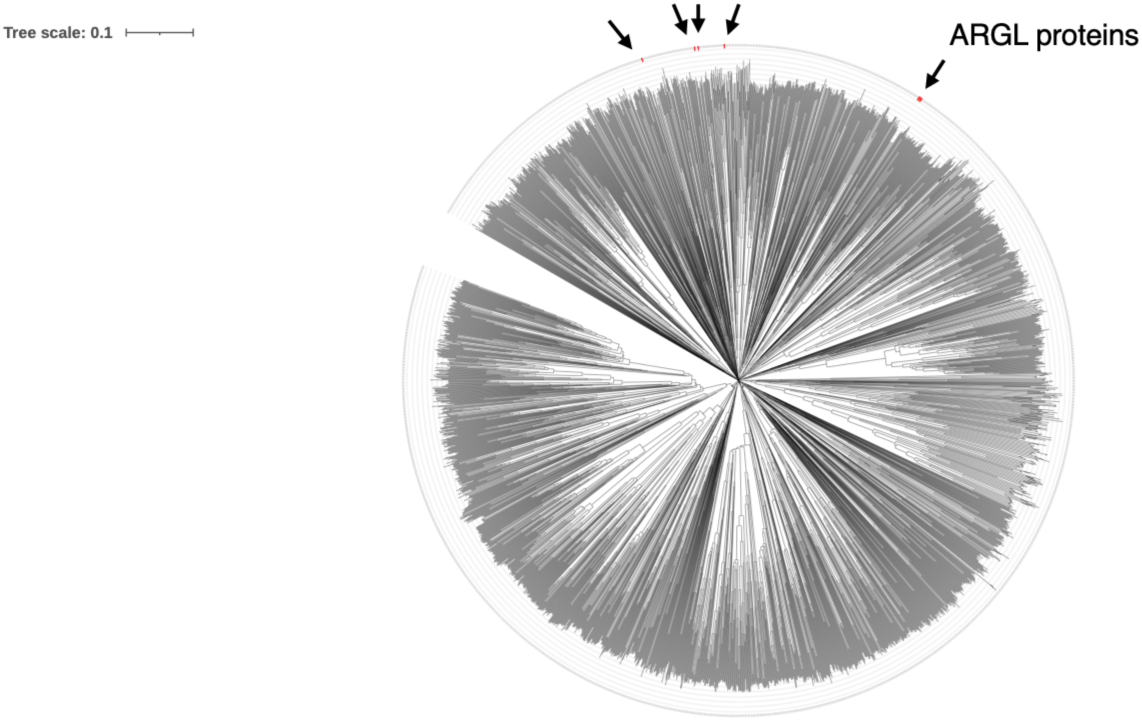
Phylogenetic tree of all 2,743 predicted *D. vitifoliae* effector proteins based on multiple sequence alignment (MSA). ARGL proteins are highlighted in red with black arrows.

**Figure S7:**
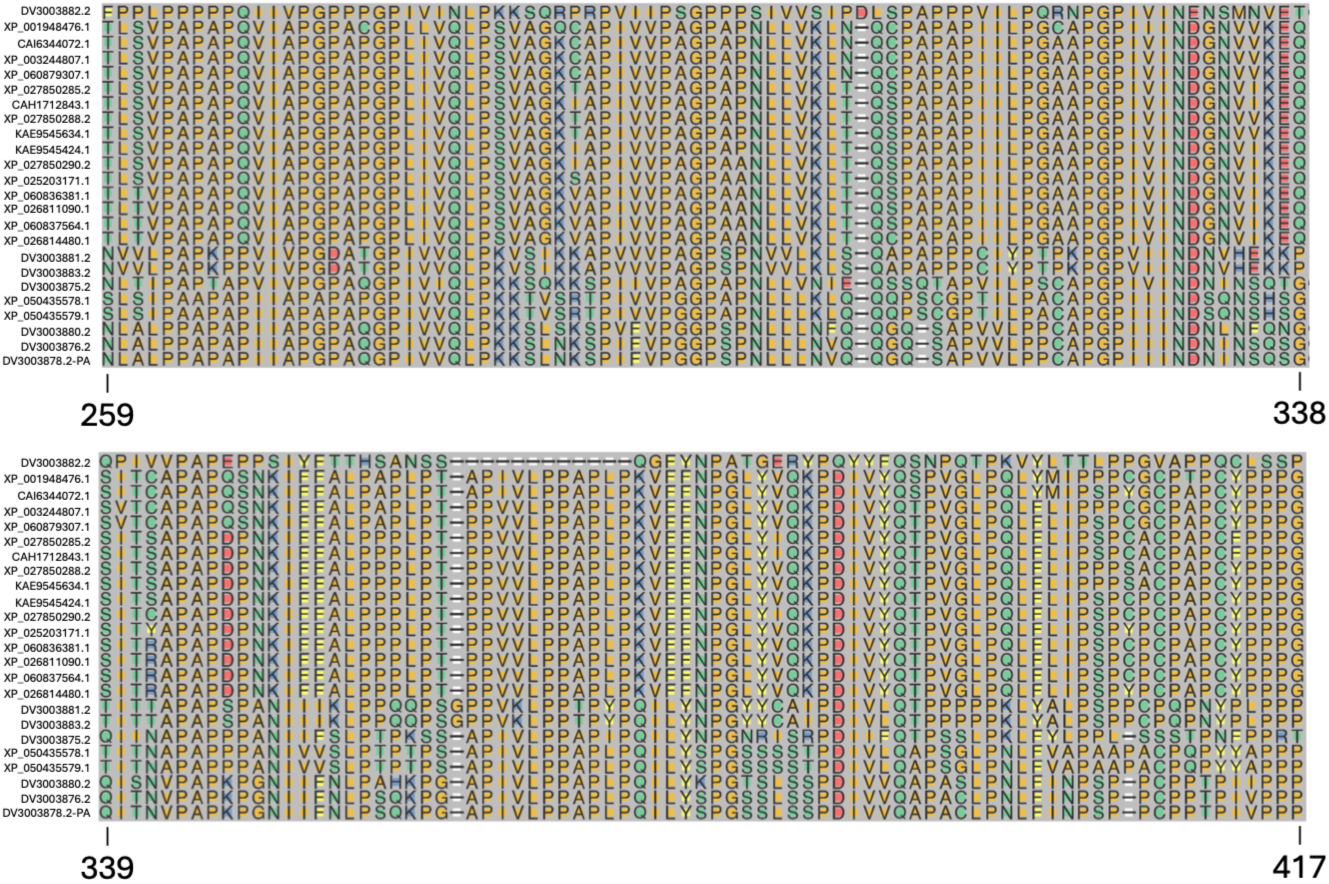
The most conserved region of ARGL proteins. A multiple sequence alignment (MSA) of ARGL proteins was performed using all seven *D. vitifoliae* predicted effector proteins at the ARGL locus and the top seventeen orthologs of ARGL protein DV3003875. Amino acid residues in the 259 to 417 positions are shown, with gene ID (*D. vitifoliae* genes) or accession number on the left.

**Figure S8:**
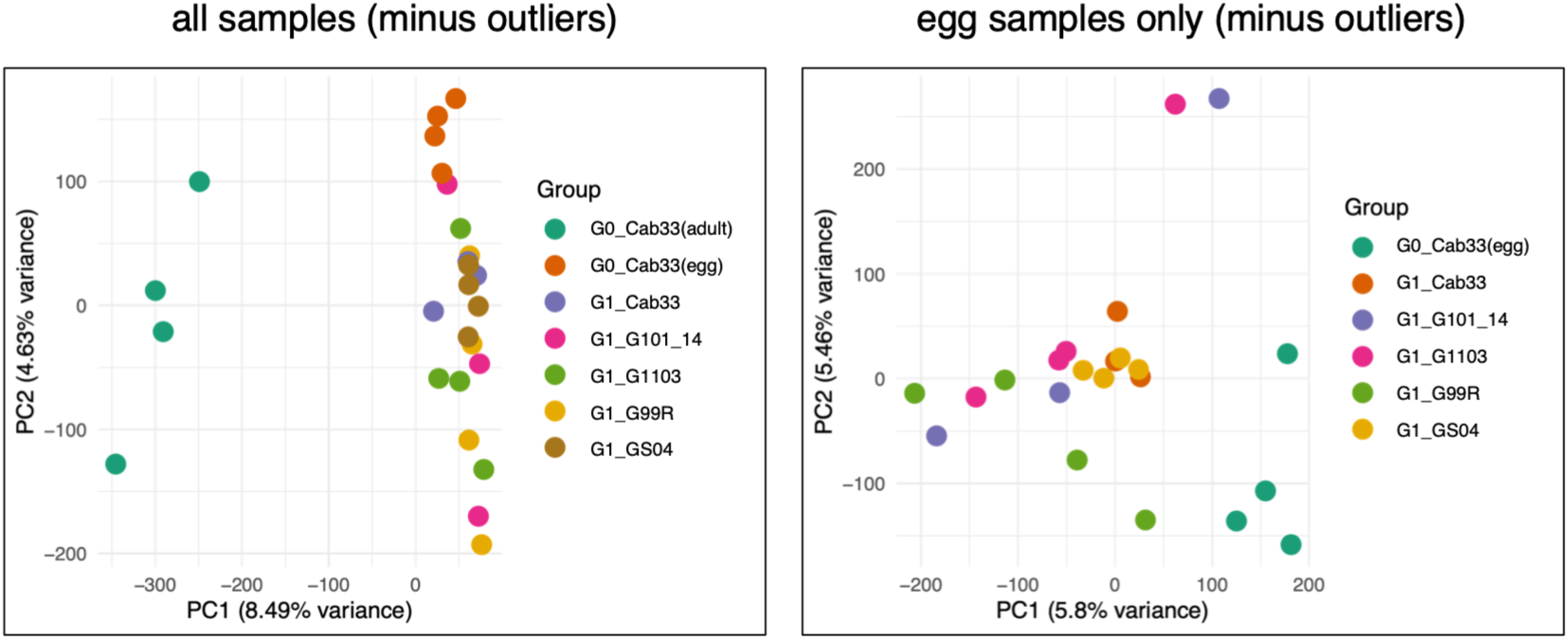
Effect of host genotype on *D. vitifoliae* CpG methylation. Principal component analysis (PCA) based on methylation data for 201,448 CpG sites for all samples minus outliers (left) and only egg samples minus outliers (right).

**Figure S9:**
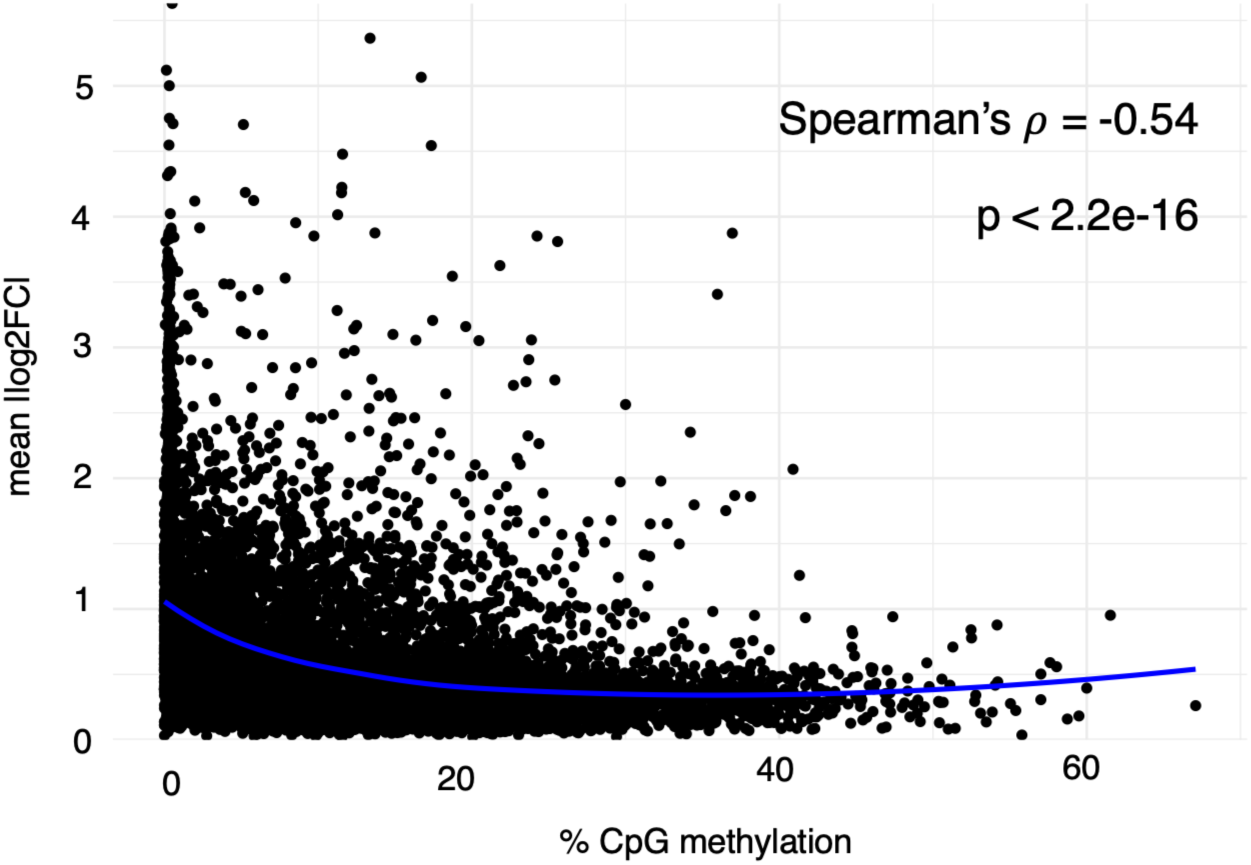
Correlation between transcriptional plasticity and gene body CpG methylation. The scatter plot shows transcriptional plasticity (mean absolute value of log2-fold-change; vertical axis) and percent CpG methylation (horizontal axis) for genes which are expressed and have CpG methylation data (n=12,368). Results of Spearman’s rank correlation test are printed in the top right of the plot. A Locally Estimated Scatterplot Smoothing (LOESS) line was fitted to the data (blue line).

## Notes

### Competing Interest Statement

The authors have declared no competing interest.

### Summary of Updates

Several typos have been corrected, included missing citation information and duplicated citations, and typos such as 'adaptative' replaced with 'adaptive'.

